# Low-usage splice junctions underpin immune-mediated disease risk

**DOI:** 10.1101/2023.05.29.542728

**Authors:** Omar El Garwany, Nikolaos I Panousis, Andrew Knights, Natsuhiko Kumasaka, Maria Imaz, Lorena Boquete Vilarino, Anthi Tsingene, Alice Barnett, Celine Gomez, Daniel J Gaffney, Carl A. Anderson

## Abstract

The majority of immune-mediated disease (IMD) risk loci are located in non-coding regions of the genome, making it difficult to decipher their functional effects. To assess the extent to which alternative splicing contributes to IMD risk, we mapped genetic variants associated with alternative splicing (splicing quantitative trait loci or sQTL) in macrophages exposed to 24 cellular conditions. We found that genes involved in innate immune response pathways undergo extensive differential splicing in response to stimulation and detected significant sQTL effects for 5,734 genes across all conditions. We colocalised sQTL signals for over 700 genes with IMD-associated risk loci from 21 IMDs with high confidence (PP4 ≥ 0.75). Approximately half of the colocalisations implicate lowly-used splice junctions (mean usage ratio < 0.1). Finally, we demonstrate how an inflammatory bowel disease (IBD) risk allele increases the usage of a lowly-used isoform of PTPN2, a negative regulator of inflammation. Together, our findings highlight the role alternative splicing plays in IMD risk, and suggest that lowly-used splicing events significantly contribute to complex disease risk.

## Introduction

Genome wide association studies (GWAS) have uncovered thousands of genetic loci associated with susceptibility to immune-mediated diseases (IMD). Over 90% of these loci are located in non-coding regions of the genome^1^, making it difficult to gain insights into causal disease biology. These non-coding disease-associated loci are enriched in gene regulatory regions, and are therefore thought to modulate gene expression^2^. Expression quantitative trait loci (eQTL) mapping has been widely used to characterise the downstream effects of genetic variants on gene expression^3, 4^. Despite the increasing number of available eQTL datasets^5^, IMD-associated loci have remained largely unexplained by existing QTL maps. For example, <u>Chun et al. 2017</u> (ref: 6) found that only 25% of IMD-associated loci colocalised with eQTLs from three immune cell types.

Multiple explanations have been put forward to justify the incomplete overlap between GWAS loci and existing eQTL maps, including a need for more diverse molecular QTL maps across disease-relevant cell types and environmental conditions. Moreover, it has recently been suggested that common variants driving complex diseases and gene expression are systematically different, and that alternative molecular QTLs (such as those affecting splicing, chromatin accessibility and chromatin interactions) may be more likely to colocalise with disease associated loci^7^. Unfortunately, most QTL mapping studies have focussed on associating genetic variation with overall levels of gene expression without considering, for example, variation in transcript isoforms. The few studies that have mapped genetic variants associated with alternative splicing (splicing quantitative trait loci or sQTLs) have shown their promise for understanding disease^8, 9^. There is thus an urgent need for sQTL maps to be constructed across a broad range of environmental contexts for disease relevant cell types.

Here, we mapped sQTLs in iPSC-derived macrophages in 12 different cellular conditions obtained from 209 individuals at two timepoints after stimulation. We quantify the extent to which alternative splicing responds to macrophage stimulation, and how sQTLs and response sQTLs (re-sQTLs) are shared across environmental contexts. We also explore the contribution of alternative splicing and sQTLs to IMD risk. Finally, we contextualise our findings within an ongoing scientific debate about the functional and evolutionary relevance of low-usage splicing events and discuss the implications of our work on the design of future transcriptomics studies.

## Results

### Macrophage stimulation initiates an alternative splicing programme in innate immune response genes

Induced pluripotent stem cell lines (iPSC) from 209 healthy unrelated individuals, generated as part of the HipSci project^10^, were differentiated into iPSC-derived macrophages (experimental protocol described in Supplementary note in Panousis et al. 2023; in press). RNA was harvested from macrophage precursors at day 0 (Prec_D0) and day 2 (Prec_D2). Naïve macrophages were exposed to a panel of 10 stimuli, and RNA was obtained and processed 6 and 24 hours after stimulation (in addition to unstimulated controls; Ctrl_6 and Ctrl_24), resulting in a total of 24 different conditions. We quantified alternative splicing from split reads (reads mapping across two splice junctions) using Leafcutter, which quantifies alternative splicing as intron usage ratios^11^. We derived an intron usage ratio matrix for each of the 24 conditions and used this for differential splicing analysis and sQTL mapping.

Macrophage precursors (Prec_D0) clustered separately from all other conditions in a UMAP projection of intron usage ratios across conditions (Figure 2a,b) (Pearson correlation coefficient between Prec_D0/Prec_D2=0.74; Supplementary Figure S1). Precursors at day 2 (Prec_D2) clustered together with the fully differentiated conditions, suggesting that macrophage splicing programmes are activated early in the seven day long differentiation process (Supplementary note in Panousis et al. 2023; in press) (Pearson correlation coefficient between Prec_D2/Ctrl_6=0.88 and Prec_D2/Ctrl_24=0.92; Supplementary Figure S1). We also observed a clear separation between cells harvested after 6 and 24 hours, an effect that was not clearly detected for unstimulated macrophages (Ctrl_6 and Ctrl_24; Figure 2a,b) . These findings demonstrate that both differentiation of iPSCs into macrophages and macrophage stimulation initiate profound and temporal alternative splicing changes.

**Figure 1:**
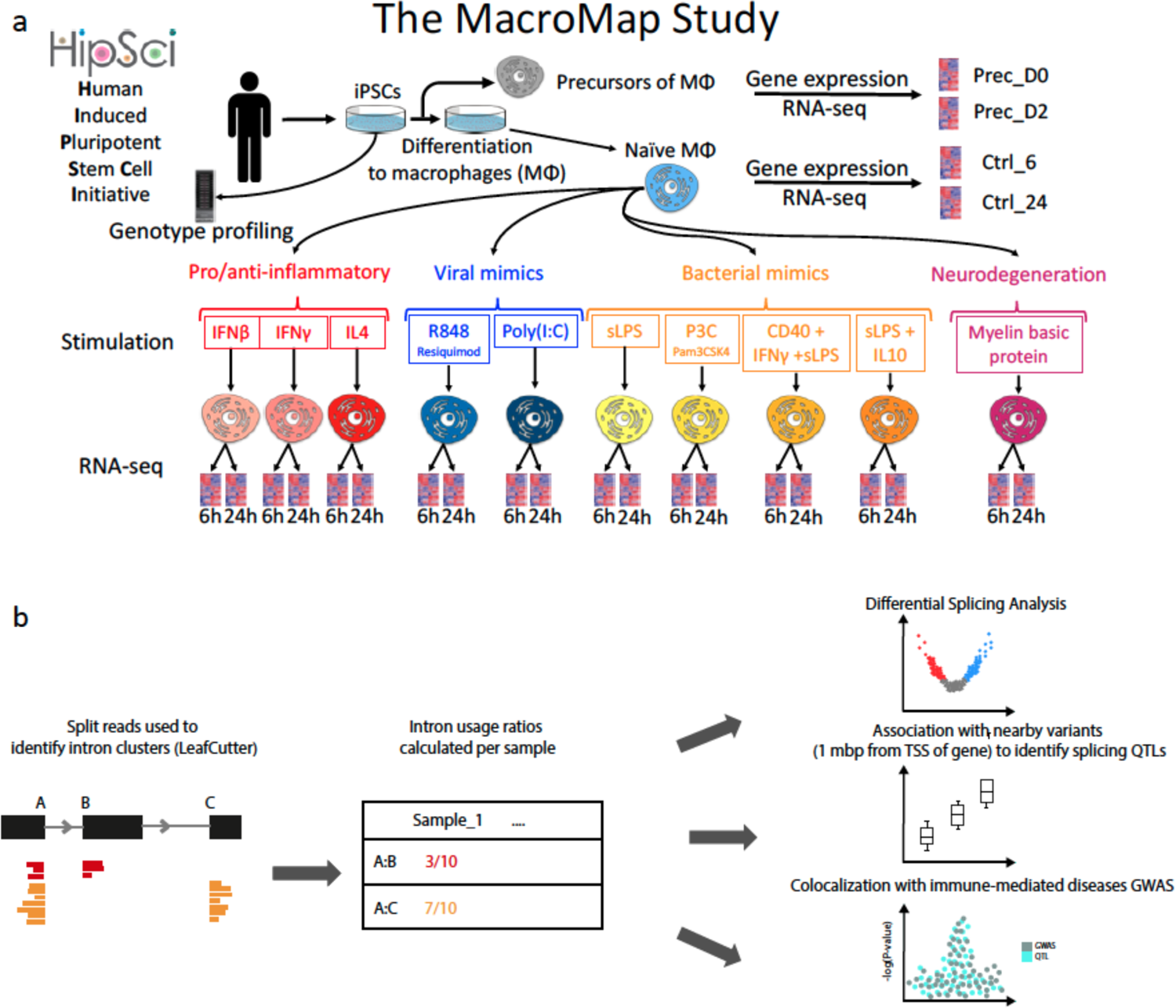
Overview of study: (a) Genotyped iPSC cell lines were differentiated into macrophages, and RNA was harvested before differentiation (Prec_D0) and 2 days after starting differentiation (Prec_D2). RNA was also harvested from differentiated macrophages at 6 and 24 hours (Ctrl_6 and Ctrl_24). Naive macrophages were then exposed to a panel of 10 stimuli and RNA was harvested at 6 and 24 hours after stimulation. (b) Split reads were used to quantify intron usage ratios on an individual level using LeafCutter. Split reads were then used for differential splicing analysis between naive and stimulated conditions, and as a quantitative trait to map splicing quantitative trait loci (sQTLs). sQTLs were then colocalised with 21 immune-mediated disease GWAS summary statistics.

**Figure 2:**
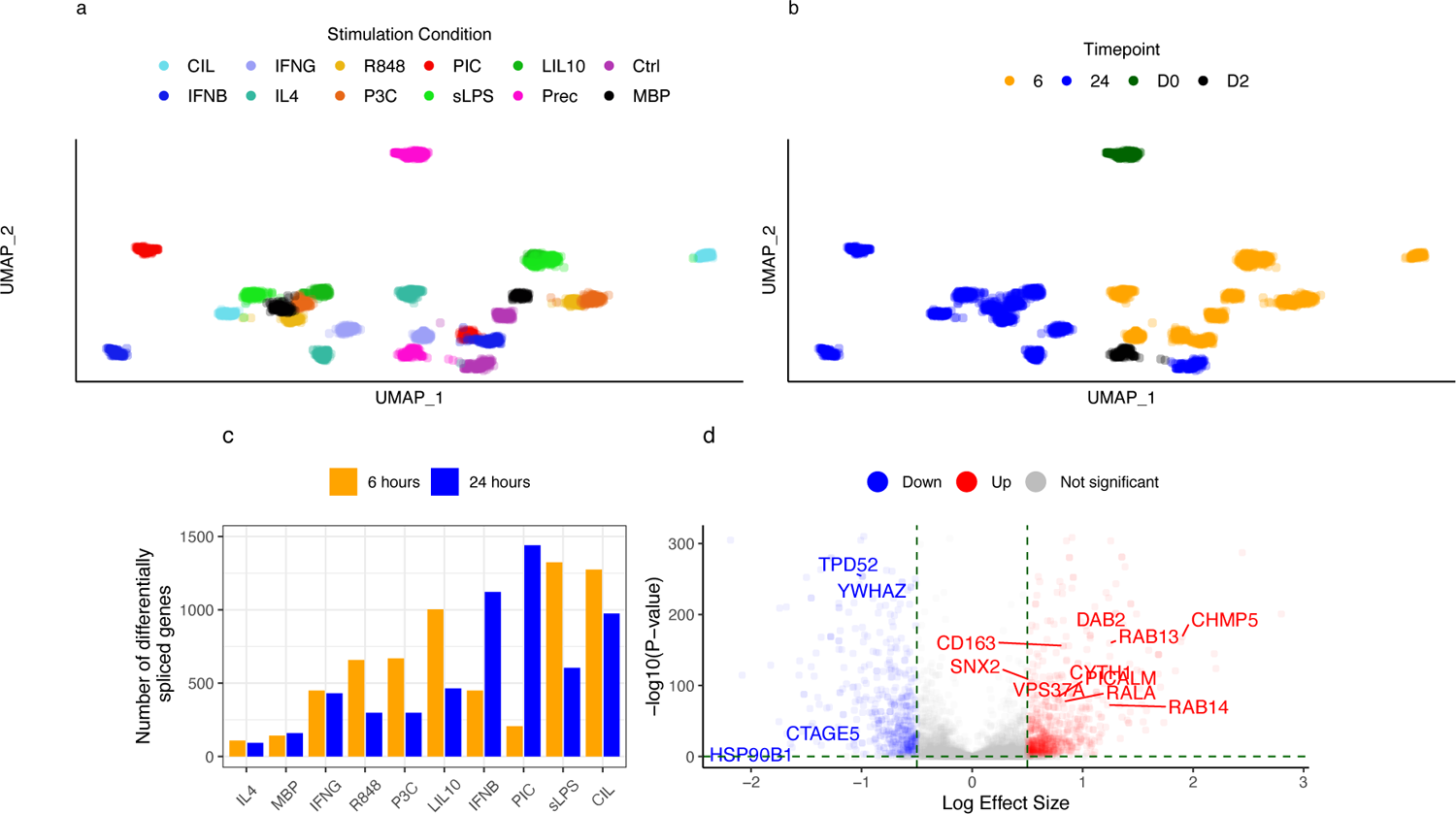
UMAP of intron usage ratios in different stimulation conditions, coloured both by (a) different stimulation conditions and (b) by time point. (c) Number of differentially spliced genes between naïve and stimulated macrophages after 6 hours (yellow) and 24 hours (blue) (d) Volcano plot showing differentially spliced genes 6 hours after sLPS stimulation, with log effect size on the x-axis (for each gene the intron with the largest absolute effect size is shown) and -log_10_ of adjusted P-value on the y-axis. Colours indicate the direction of intron usage change (blue indicating reduced usage and red indicating greater usage in stimulated cells versus naïve cells). Genes that belong to the “vesicle-mediated transport” REACTOME pathway are indicated (not all genes are shown for clarity).

To identify genes that are differentially spliced upon immune stimulation, we undertook differential splicing analysis using LeafCutter (Methods). We identified a total of 3,464 genes with altered splicing in stimulated versus unstimulated macrophages after either 6 or 24 hours (adjusted P-value < 0.05 and |log effect size| > 0.5). Macrophages stimulated with IL4 had the fewest differentially spliced genes (110 and 94 genes after 6 and 24 hours, respectively; Figure 2c), in line with previous reports that showed stimulating macrophages with IL4 had little impact on splicing^12^.

We next undertook pathway enrichment analysis to identify REACTOME pathways enriched with differential spliced genes following stimulation. Six hours after bacterial stimulation with LPS (sLPS_6), differentially spliced genes were enriched for pathways relevant to macrophage response, including “Cytokine Signalling” (84 genes; P-value=9.1×10^-8^), “Vesicle-mediated transport” (68 genes; P-value=10^-6^) and “Class I MHC mediated antigen processing and presentation” (47 genes; P-value=7.95×10^-6^) (Supplementary Table S1). Several genes coding for members of the RAB family of proteins were enriched within the vesicle-mediated transport pathway, which is also known to be regulated via alternative splicing^13^ (including *RAB1B*, *RAB4A*, *RAB9A*, *RAB11A*, *RAB13*, *RAB14* and *RABEPK;* Figure 2d, Supplementary Table S1). RABs are GTPases that regulate membrane trafficking by controlling the formation, movement and recycling of vesicles^14^. For example, we observed greater usage of the first exon of a non-canonical transcript of *RAB13* (RAB13-205), coupled with lower usage of the canonical first exon (RAB13-201) (difference in Percentage Spliced In (**Δ**PSI)=0.046 and −0.05, respectively). The two annotated transcripts RAB13-201 and RAB13-205 result in protein products of 203 and 122 amino acids respectively, which indicates that a different *RAB13* protein isoform is produced following stimulation with LPS.

Our stimulation panel also included viral stimuli, which enabled us to explore if viral response genes were differentially spliced upon stimulation. We observed that six hours after stimulation with dsRNA viral mimic PolyI:C (PIC_6), genes involved in viral sensing and RIG-I/MDA5-mediated activation of the antiviral cytokine Interferon β (IFNβ) are differentially spliced (Supplementary Table S1). RIG-I/MDA5 belong to the RIG-I-like family of receptors which sense dsRNA and activate type I interferons in response (e.g. IFNβ)^15, 16^. Differentially spliced genes include genes coding for the RIG-I and MDA5 receptors themselves (*DDX58* and *IFIH1* respectively), but also genes involved in IFNβ regulation, such as *TRIM25*^17^, *TANK*^18^, and *RIPK1*^19^. These findings are supported by previous work showing that regulatory components of the RIG-I/MDA5-mediated IFNβ activation pathway have different isoforms that modify antiviral response^20–22^.

Our differential splicing analysis reinforces the role of alternative splicing in regulating important innate immunity processes (e.g. vesicle-mediated transport and viral sensing and response), but also highlights the specificity of splicing junction usage in response to specific stimuli. Although the role of alternative splicing in response to stimuli has been previously reported^23^, our comprehensive and well-powered screen captures pathway-level changes in splice junction usage in response to a broad range of stimuli. Furthermore, these findings demonstrate that iPSC derived macrophages can faithfully recapitulate known biological pathways activated by different stimuli. This motivated us to investigate how genetic variation affects alternative splicing in macrophages and how such genetic variation may, in turn, predispose to IMDs.

### Macrophage stimulation increases the number of genes with significant sQTL effects

To understand which alternative splicing events were affected by genetic variation, we mapped splicing QTLs using intron usage ratios as a quantitative trait. Intron usage ratios were normalised (quantile normalised and rank-based inverse-normal transformed) and introns with low variance (standard deviation < 0.005), or intron clusters without split reads in more than 40% of samples, were excluded (Methods). We identified a median of 82,058 introns (75,987-105,841 introns across conditions with the greatest number of introns seen in Prec_D0; Supplementary Figure S2). Each gene had a median of 7 introns and up to 10,851 genes were quantified per condition.

We mapped sQTLs within a ± 1 Mbp window centred around the transcription start site (TSS) of each gene. We also used multivariate adaptive shrinkage (mash^24^) to compare sQTL effect sizes between naïve and stimulated conditions (we used Ctrl_24 as our baseline condition). Mash reports a significance measure known as local false sign rate (LFSR; Methods), which we used to identify sQTLs whose effect sizes change significantly upon stimulation (response sQTLs).

We called significant sQTLs at a false discovery rate (FDR) < 0.05. Across all conditions we detected a total of 5,734 sGenes (median number of sGenes per condition=1580 and Prec_D2 had the most sGenes=1,881) (Figure 3a,b). Of these, 878 sGenes (15.3%) had at least one response sQTL (LFSR < 0.05). We then asked which of the two stimulation timepoints (6 and 24 hours) was more likely to have response sQTL effects across stimulation conditions. Up to 29% of response sGenes per condition had a response sQTL after 24 hours that was not detected after 6 hours. Conversely, up to 72% of response sGenes per condition had a response sQTL after 6 hours that was undetectable after 24 hours. This suggests that response sQTLs are more likely to be detected 6 hours after stimulation (Figure 3c), in agreement with previous work that suggested 4-6 hours for optimal detection of transcriptomic changes following macrophage stimulation^25, 26^.

**Figure 3:**
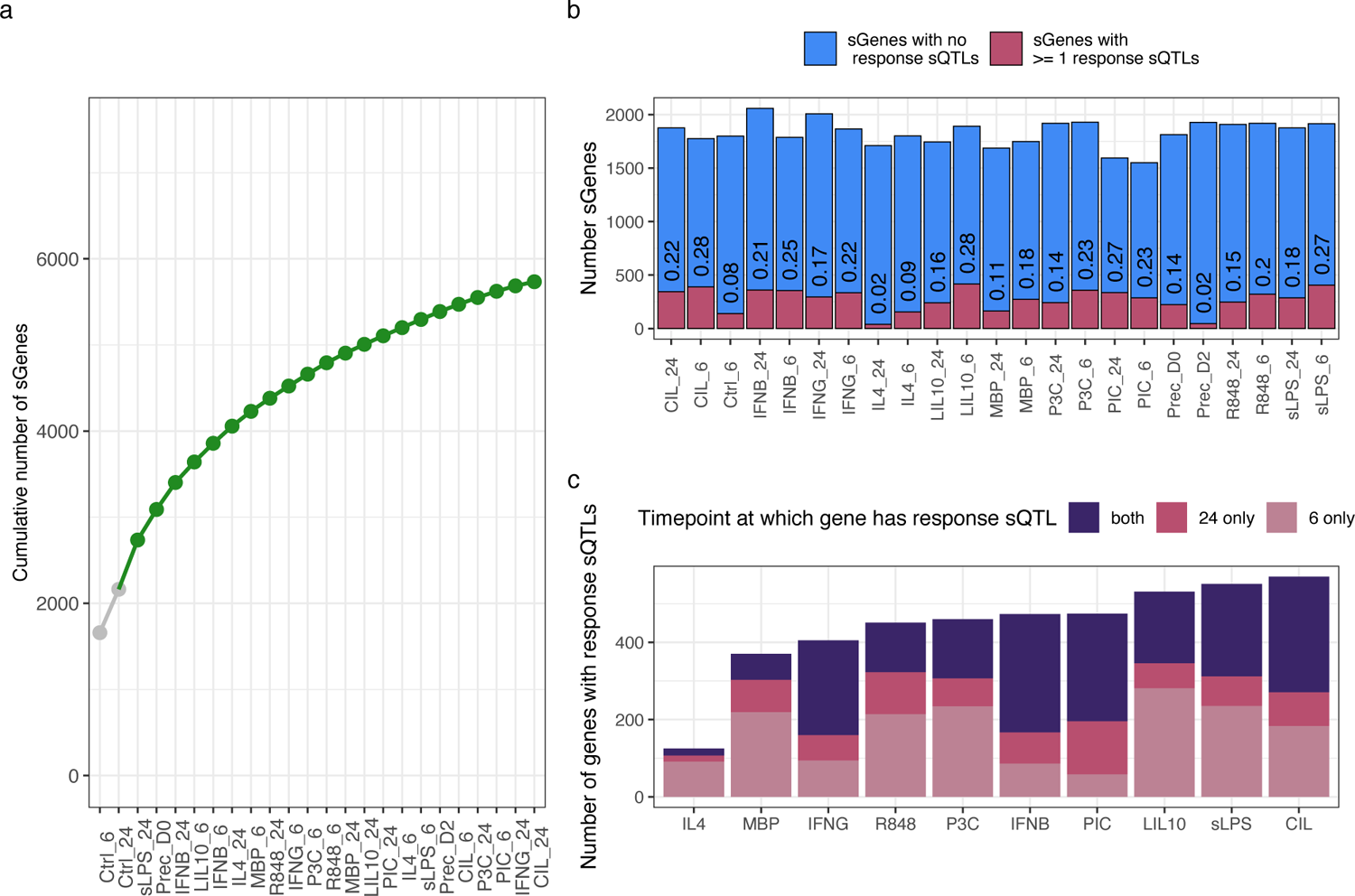
(a) cumulative number of genes with significant splicing QTL effects, with unstimulated conditions indicated in grey (b) Total number of significant sQTLs per condition and proportion of response sQTLs within each condition (sQTLs with LFSR < 0.05; Methods). (c) Number of genes, per condition, with at least one response sQTL at 6 hours, 24 hours or both.

Similar to previous reports^27^, we found that lead sQTL SNPs are located closer to intron boundaries than to the TSS of their genes. On average per condition, 24.5% of sGenes had a lead SNP within 10 kbp of their TSS, compared to 46.8% within 10 kbp of either the 5’ or 3’ intron boundaries (Supplementary Figure S3). Previous sQTL mapping efforts have also shown that sQTL signals are largely independent from eQTL signals, and therefore overall levels of gene expression and alternative splicing are genetically regulated via distinct mechanisms^9, 28, 29^. To verify this, we performed statistical colocalisation (Methods; ref 30) between sQTLs and eQTLs derived from the same data (Panousis et. al. 2023; in press). In line with previous findings, we found that on average across conditions, only 25% of sGenes might share a single causal variant with an eGene in the same condition (PP4 ≥ 0.1; Supplementary Figure S4; Methods), indicating that the majority of sQTL signals are largely independent from eQTL signals. We therefore hypothesised that colocalisation of sQTLs with disease-associated loci may identify additional disease effector genes distinct from those already implicated via eQTLs.

### Splicing QTLs identify additional GWAS loci effector genes not detected by expression QTLs

Macrophage sQTL maps can also be used to build hypotheses about how IMD risk loci confer risk by dysregulating alternative splicing. To this end, we used statistical colocalisation (using coloc; Methods) between sQTL and GWAS association signals to quantify the probability of a sQTL sharing a causal variant with genetic association signals from 21 IMD GWAS summary statistics (downloaded from GWAS catalogue^31^; Methods and Supplementary Table S2).

Across all 21 IMDs, we identified 707 unique genes (1,337 introns) with an sQTL signal in at least one condition that likely shares a causal variant with an IMD risk locus (PP4 ≥ 0.75; Figure 4a). Moreover, for 68 (9.6%) of these genes, the risk locus colocalised with a response sQTL, indicating that the genetic effects of these variants on alternative splicing could not be detected in unstimulated macrophages. Based on hierarchical clustering of LFSR values, Prec_D2, IL4_6 and IL4_24 had the fewest response sQTLs that colocalised with GWAS signals, while sLPS_6, CIL_6 and LIL10_6 (all stimulated with LPS) yielded the most, recapitulating results from the differential splicing analysis (Figure 4b).

**Figure 4:**
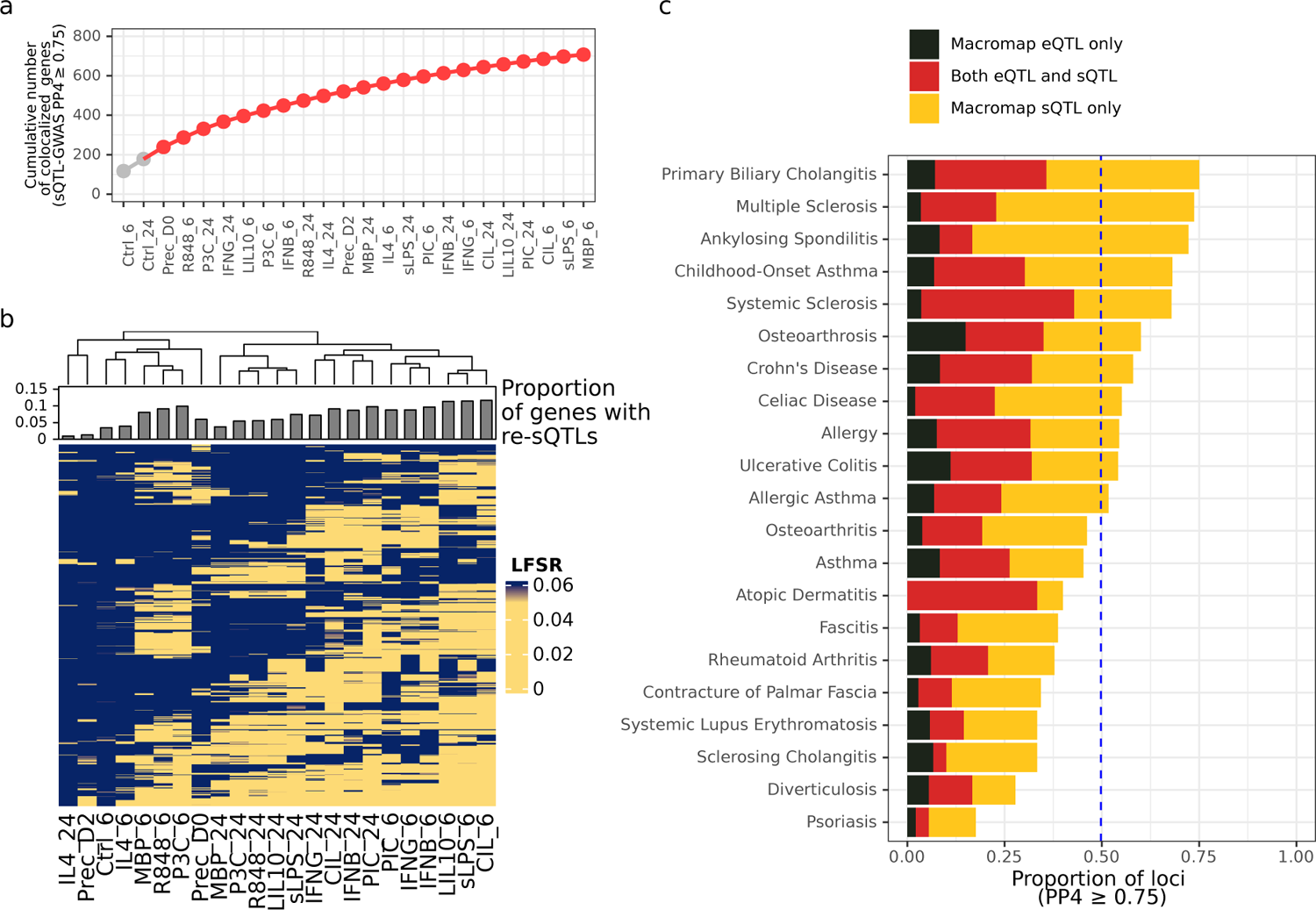
(a) Cumulative number of genes with a GWAS-sQTL (PP4 ≥ 0.75) across different conditions, with unstimulated conditions shown in grey on the left (b) Heatmap and hierarchical clustering of LFSR values for all the colocalised sQTL effects (PP4 ≥ 0.75) across all 21 IMDs. On top of the heatmap is a barplot showing the proportion of colocalised sQTL effects that are response sQTLs (LFSR < 0.05) (c) Proportion of genome-wide significant loci that share a single causal variant (PP4 ≥ 0.75) with an eQTL only, an sQTL only or both.

We then compared how many tested loci colocalised with each type of molecular QTL and found that 50.4% (771/1,528) of tested loci were likely to share a single causal variant with either an eQTL, sQTL or both. Recently, <u>Mountjoy et al. 2021</u> (ref: 32) colocalised 50.7% of tested GWAS loci with protein and expression QTLs from 92 tissues and cell types. In comparison, our high colocalisation yield from a single cell type shows the promise of profiling the transcriptome of a relevant cell type in understanding the effects of GWAS loci. Moreover, unlike other tissue-level QTL maps, interpreting these colocalisations in the context of macrophages potentially aids the interpretation and functional follow-up of these loci.

Approximately half of the colocalised loci (385 loci or 25.2% of tested loci) colocalised solely with an sQTL (sQTL PP4 ≥ 0.75 > eQTL PP4), clearly demonstrating both the value of sQTLs for identifying GWAS effector genes and the important role that alternative splicing plays in complex disease risk (Figure 4c).

### Lowly-used alternative splicing events underlie complex disease risk

We next sought to characterise the colocalised sQTL introns, by asking how often colocalised sQTL splicing events are used in observed transcripts. There is ample evidence that lowly-observed aberrant splicing underpins several inherited diseases such as Spinal Muscular Atrophy and Duchenne Muscular Dystrophy^33–35^, but it is unclear to what extent lowly-observed splicing events contribute to complex diseases.

We observed that 53.4% of colocalised sQTL introns have a mean intron usage ratio (IUR) < 0.1 across samples (Figure 5a). Over 96% of these introns had non-zero usage in at least 30 samples (Figure 6c), indicating that these splicing events can be reliably observed in multiple RNA-seq samples and individuals, but with relatively low IUR. Moreover, 50.6% of these introns are not found in any annotated transcripts in GENCODE v27, whereas only 12% of introns with mean IUR ≥ 0.1 are absent from GENCODE v27 (Figure 6b), in line with previous reports showing that lowly-observed splicing events tend to be unannotated in transcript databases^36, 37^.

**Figure 5:**
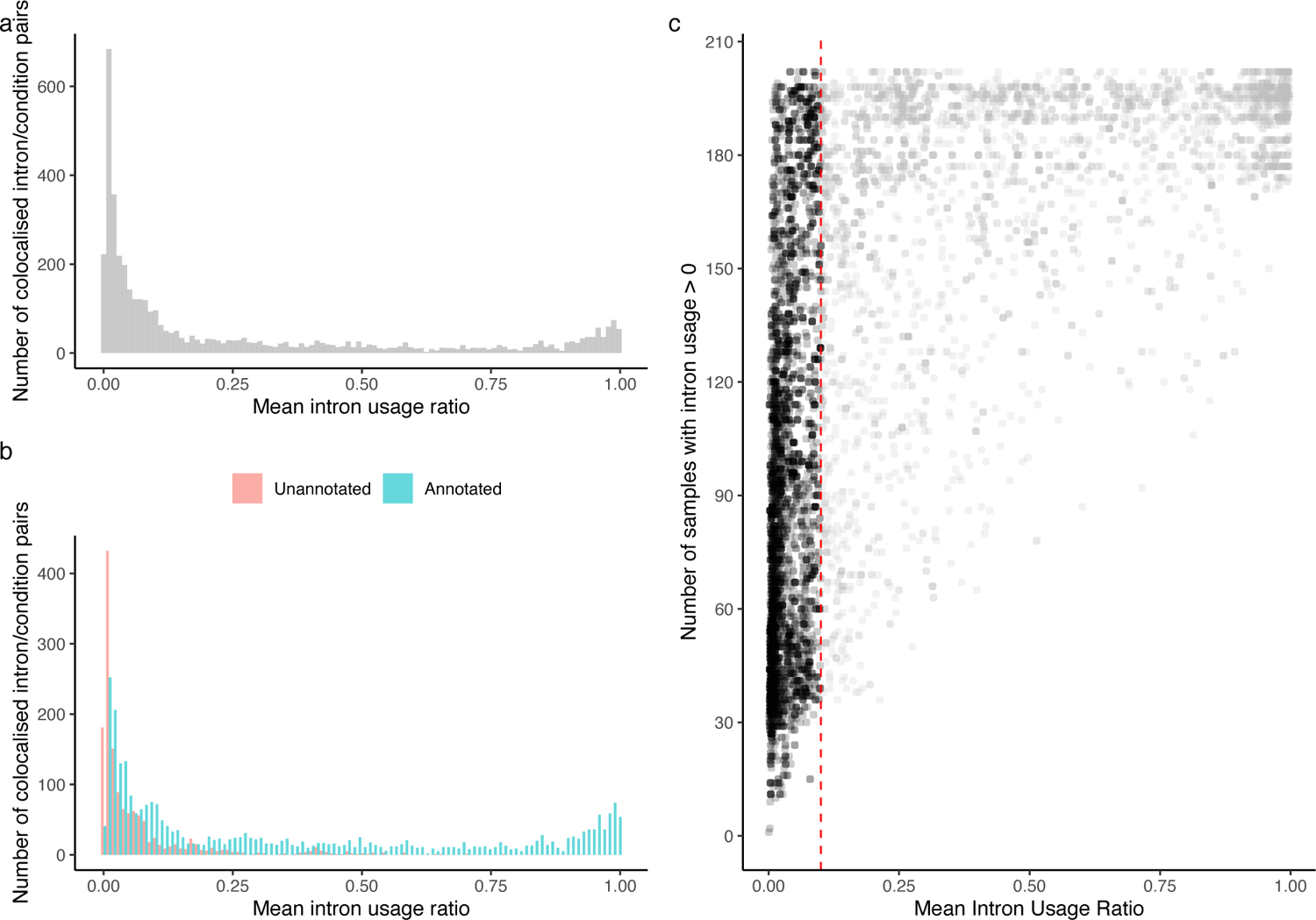
(a) distribution of mean intron usage ratio for colocalised introns, showing a peak close to 0, and (b) coloured by annotation in GENCODE v27, showing an enrichment of unannotated introns among introns with low mean usage ratio (c) number of samples in which the intron is supported by at least one split read (y-axis) shown against the mean intron usage ratio of each sample (x-axis). Red vertical line at mean intron usage ratio = 0.1.

**Figure 6:**
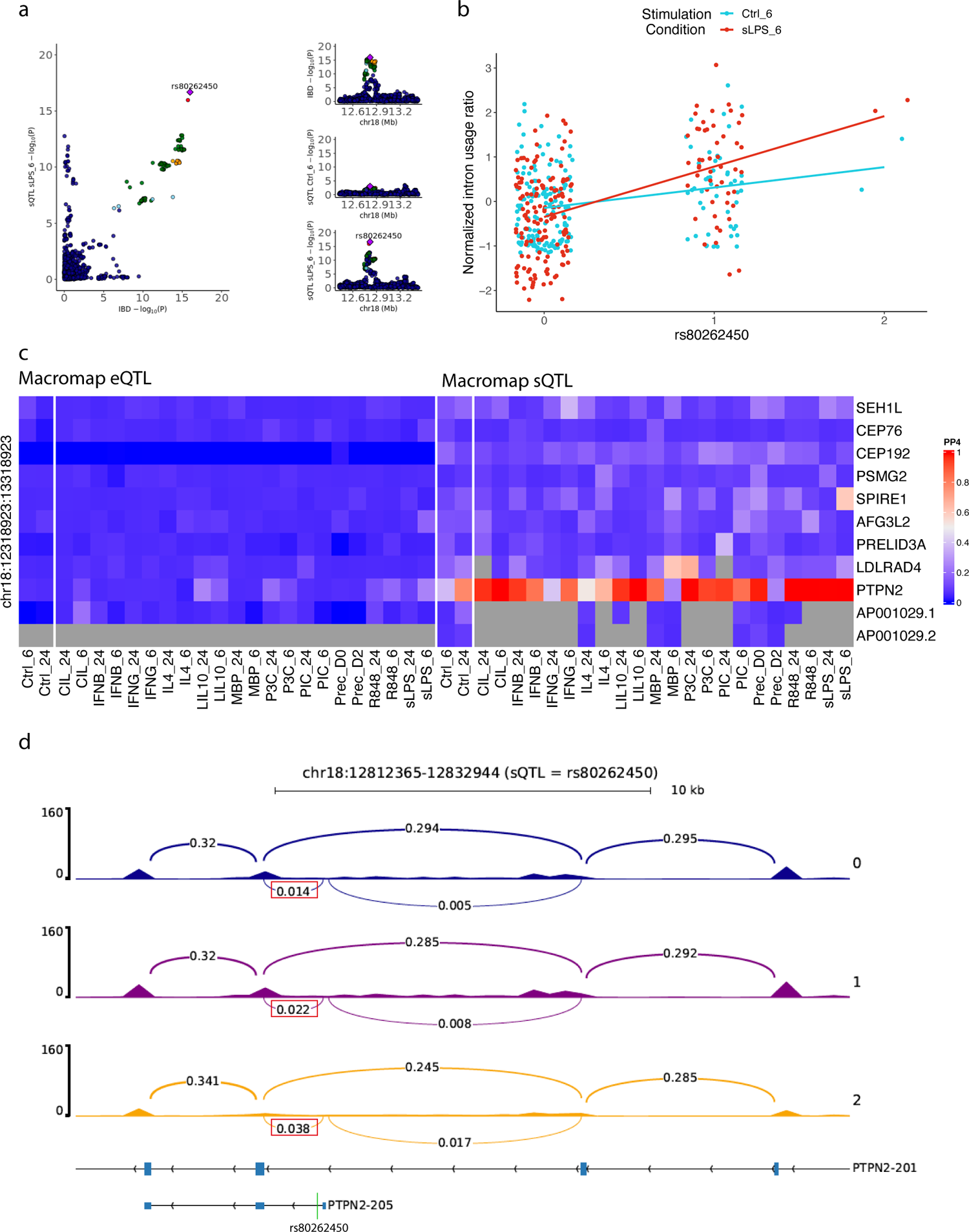
Example of colocalisation between an IBD risk locus at 18p11.21 and an sQTL for *PTPN2.* (a) Regional association plots of the IBD association signal, and sQTL association signal in unstimulated macrophages (Ctrl_6) and macrophages stimulated with sLPS after 6 hours (sLPS_6), (b) Normalised intron usage ratios of different genotypes of the lead IBD SNP rs80262450 in Ctrl_6 and sLPS_6, (c) Heatmap showing evidence of colocalisation (PP4) between the IBD association signal at 18p11.21 and all macrophage eQTLs/sQTLs in the locus (in all conditions; PP4 for top splice junction is shown per gene), (d) RNA-seq coverage of the intron cluster where the *PTPN2* sQTL effect is detected in sLPS_6. Bars represent the number of reads and arcs represent the usage of different splice junctions (only five splice junction ratios are shown for clarity and the colocalised sQTL splice junction is indicated in a red box). Canonical transcript PTPN2-201 and non-canonical transcript PTPN2-205 are shown underneath, with blue boxes representing introns and the position of rs80262450 within PTPN2-205 is shown by the green line.

Our observation that over half of colocalised sQTL introns are lowly used and that they tend to be unannotated strongly emphasises the need to investigate their functions and role in the context of complex disease. For example: are low-abundance isoforms translated into protein products or do they exert gene regulatory functions? What is the effect of up- or down-regulating these isoforms on different cellular phenotypes? Sampling these rare transcripts, an avenue that has remained largely unexplored in most large-scale transcriptomic cohorts, will thus shed light on the transcriptomic effects of IMD-associated risk loci.

### A rare alternative splicing event likely underpins inflammatory bowel disease risk at the *PTPN2* locus

To demonstrate how sQTLs for lowly-used introns can dysregulate alternative splicing and predispose to IMDs, we further investigated a *PTPN2* sQTL that implicates a lowly-used intron that colocalised with an inflammatory bowel disease (IBD) associated risk locus at 18p11.21. Multiple lines of evidence, including coding variants associated with monogenic IBD^38, 39^ and mouse knock-out models^40, 41^, have suggested *PTPN2* is the effector gene at 18p11.21, though this remains to be established. Moreover, It is not yet known if and how common IBD-associated SNPs affect the expression of *PTPN2*.

We observed that the lead IBD SNP at 18p11.21 (rs80626450; 18-12818923-G-A) is associated with higher risk of IBD and with increased usage of intron 1-2 of the non-canonical transcript PTPN2-205 (chr18:12,817,365-chr18:12,818,944; Figure 6b,6d). rs80626450 is located 21 base pairs downstream of the donor splice site of exon 1 of PTPN2-205, and is the lead SNP for both the sQTL and IBD association signals, strongly suggesting its involvement in the aberrant splicing event at this locus. The PTPN2 sQTL signal colocalised (PP4 ≥ 0.9) with the IBD signal in 13 conditions, but did not colocalise with any eQTLs mapped from the same data (eQTLs mapped in Panousis et. al. 2023; in press). Despite the relative rarity of this transcript, we detected this colocalisation using GTEx sQTL summary statistics^10^, where this sQTL signal was also colocalised with the IBD locus (PP4 ≥ 0.9) in 14 tissues, including whole blood (Supplementary Table S3 and Supplementary Figure S5), indicating that this rare splicing event can be reliably detected in a large number of tissues from an independent dataset.

The directions of effects of rs80626450 on intron usage and IBD risk (effect size in sLPS_6=1.21771 and odds ratio=1.17, respectively) suggest that an increase in the relative abundance of PTPN2-205 is associated with increased risk of IBD. Given that mouse knock-out studies of *PTPN2* suggest the gene plays an anti-inflammatory role in macrophages, we hypothesise that increased usage of PTPN2-205 attenuates the role of *PTPN2* as a negative regulator of inflammation, which in turn increases the risk of IBD.

### sQTL colocalisations converge on dysregulated pathways in IMDs

For example, we found that two IBD-associated loci colocalised with sQTL signals for genes that interact with the RAB GTPase Rab35. Rab GTPases are a diverse set of molecules that regulate different aspects of vesicle-mediated transport, with over 70 RAB GTPAses discovered so far^42^. RAB GTPases are activated by Guanine Exchange Factors (GEF), and subsequently recruit effector proteins, including proteins required for vesicle uncoating, movement, tethering and fusion, which enables cargo trafficking across cell compartments and membranes^15^.

IBD-associated loci at 12q12 and 1q31.3 colocalise with *LRRK2* and *DENND1B* sQTLs in 5 and 21 conditions, respectively (PP4 ≥ 0.75; Supplementary Table S4 and S5; Figure 7). *DENND1B* is a GEF that activates Rab35, while Rab35 was shown to be a bona fide substrate for *LRRK2* ^43, 44^, although the exact effect *LRRK2* has on Rab35 is still debated^45–47^. Rab35 regulates endosomal recycling of both integrins and cadherins, effectively maintaining a balance between cell adhesion and cell migration. Moreover, evidence suggests that endosomal recycling in epithelial cells helps maintain apical and basolateral polarity, and that Rab35 plays a role in maintaining this polarity^48, 49^. The fact that both *LRRK2* and *DENND1B* interact with Rab35 suggests that endosomal recycling is frequently dysregulated by common IBD-associated variation.

**Figure 7:**
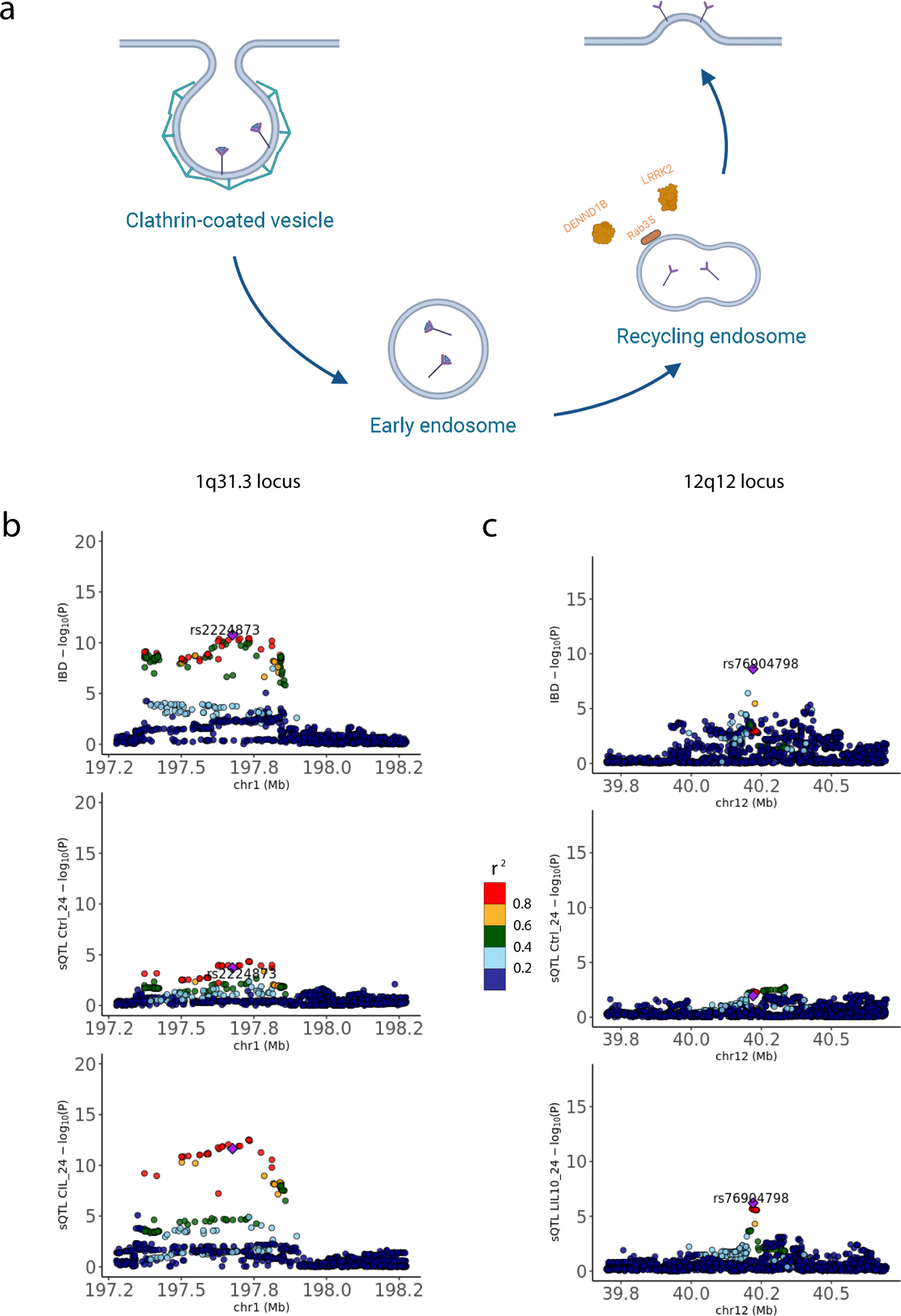
(a) Diagram showing different stages of endocytosis and endosomal recycling. Rab35 and two of its interactors *LRRK2* and *DENND1B* are shown on a recycling endosome. Regional Manhattan plots of the two IBD-associated loci (created with BioRender.com) (b) 1q31.1 and *DENND1B* sQTL in naïve and stimulated macrophages (LIL10_24) and (c) 12q12 and *LRRK2* sQTL in naive and stimulated macrophages (CIL_24) showing high colocalisation evidence with the IBD-associated locus in the stimulated condition sQTL, but not in the naïve condition sQTL. IMD-sQTL colocalisations in disease-relevant cell types can reveal how genetic variation dysregulates biological pathways that enable the cell to perform its normal functions.

Two *DENND1B* splice junctions with different donor splice sites colocalise with the IBD-associated signal 1q31.3 (chr1:197,715,074-197,772,868 and chr1:197,715,074-197,735,586). The first splice junction is located in the coding sequence of the canonical transcript of *DENND1B* (DENND1B-211), while the second is part of the 5’ untranslated region of another protein-coding transcript (DENND1B-201). Lead sQTL SNPs for both transcripts have inconsistent directions of effect in different stimulation conditions where the colocalisations are detected (Supplementary Figure S6). This suggests that the effect of the risk-increasing allele on relative transcript abundances is highly context-dependent.

Unlike *DENND1B*, all *LRRK2* sQTLs with high colocalisation probabilities (PP4 ≥ 0.75 in 5 conditions) had consistent directions of effects in the conditions where the colocalisation was detected. For example, the *LRRK2* sQTL with the highest colocalisation probability was observed in LIL10_24 (PP4=0.975), where the risk-increasing allele of the lead GWAS SNP (rs76904798; 12-40220632-C-T) decreases the usage of an *LRRK2* splice junction (chr12:40,316,403-40,319,988; beta=-0.75), and also increases the risk of IBD (beta=0.105). This indicates that decreased usage of this splice junction increases risk of IBD, and conversely that inclusion of this splice junction may have a protective effect against IBD (Supplementary Figure S6). Although we demonstrate how *LRRK2* and *DENND1B* contribute to a dysregulated endosomal recycling in IBD in different environmental contexts, more research is still needed to understand the exact mechanisms by which the isoforms of these two genes affect such an important biological process in macrophages.

## Discussion

In this work, we mapped splicing QTLs in iPSC-derived macrophages to understand how splicing is genetically controlled in macrophages under 10 different environmental contexts. We found thousands of genes with significant sQTL effects across our array of conditions, and that a considerable proportion of these genes have different effect sizes upon stimulation and were thus defined as response sQTLs.

The primary motivation behind several QTL mapping efforts is to understand the transcriptomic consequences of disease-associated genetic variation. Recently, <u>Mostafavi et</u> <u>al. 2022</u> (ref 8) suggested that eQTL and GWAS studies are powered to discover systematically different types of variants, and that this may partially explain the limited colocalisation between eQTLs and GWAS risk loci. More than half of the IMD-associated risk loci that we tested likely share a single causal variant with either an eQTL or sQTL, with sQTLs solely contributing half of these loci, which clearly demonstrates the added value of sQTLs. This echoes previous work that showed the promise of sQTLs in closing the colocalisation gap^9, 12, 50^. Along similar lines to <u>Mostafavi et al. 2022</u>, we attribute the large number of sQTL colocalisations to three important features of our discovered sQTLs that make them likely to colocalise with GWAS association signals. First, unlike eQTL variants, sQTL variants tend to be located around intron splice sites, rather than TSS of genes (Supplementary Figure S3), closer to intronic GWAS association signals that do not colocalise with eQTL signals. Indeed, 63.4% of the lead SNPs in the GWAS loci that we tested were intronic variants. Second, sQTL signals largely do not colocalise with eQTL signals, suggesting that they are driven by distinct genetic associations (Supplementary Figure S4). Third, we show that a considerable number of colocalisation events implicate unannotated introns that are lowly used across transcripts. These subtle changes in intron usage are unlikely to be reflected in overall levels of gene expression levels and may therefore remain undetected in eQTL studies.

Currently, it is unclear whether lowly-used splicing events are simply splicing errors^37, 51^ or if they have functional consequences^52, 53^. For example, <u>Pickrell et al. 2010</u> (ref 37) interpreted the lack of evolutionary conservation around lowly-used splice sites as evidence that they are functionally-irrelevant. This interpretation has been debated over the past decade^54^. Here, we show that many of these lowly-used splice junctions may underpin disease-associated genetic effects on alternative splicing, and should not be considered noisy and functionally irrelevant. The *PTPN2* example is particularly intriguing as the implicated splice junction falls within an Alu element, a class of Short Interspersed Nuclear Elements (SINE; Dfam E-value=4.7×10^-98^) that constitute 11% of the human genome^55^. RNA binding proteins normally repress the expression of newly-incorporated Alu elements, but decreased repression over long evolutionary periods provides a substrate for new functions ^56, 57^. The risk-increasing effect of the lowly-used *PTPN2* splice junction could therefore represent a harmful evolutionary by-product that attenuates the anti-inflammatory effect of *PTPN2*. This remains to be validated by functional studies that profile the functional consequences of *PTPN2* isoforms that contain this splice junction. With the recent success of RNA therapeutics such as splice-switching antisense oligonucleotides (ASO), it may be possible to “contain” these evolutionary side effects via therapeutic interventions that decrease the proportion of these lowly-used splice junctions^58, 59^. This should provide incentive for the complex disease and transcriptomics communities to understand and interrogate the functional consequences of the splicing events that underpin complex disease risk.

Although our dataset represents the largest resource for studying how alternative splicing is genetically controlled in iPSC-derived macrophages, we acknowledge two main limitations. First, although macrophages differentiated using our experimental protocol have been shown to be transcriptionally similar to monocyte-derived macrophages^60^, they still do not capture the local environment of tissue-resident macrophages. This may limit our ability to understand the effect of local micro-environments on the transcriptome of macrophage^61^. Second, intron usage ratios do not provide information about the relative abundance of full transcripts. For example, we demonstrated the potential effects of splicing events on the canonical transcript of *PTPN2* and on PTPN2-205, but it is still unclear which additional transcripts these splicing events may have originated from. Fortunately, long-read sequencing and its algorithms for isoform quantification are becoming increasingly mature^62–64^. We therefore expect that a more direct quantification of transcript usage will be attainable, and that it will make alternative splicing a more routine part of transcriptomic profiling studies.

In summary, our findings highlight an important role for alternative splicing in macrophage response to environmental cues, and that its dysregulation explains a large proportion of IMD-associated susceptibility loci. We anticipate that improved long-read sequencing technologies will facilitate whole isoform quantification in different cellular contexts, which will open the door for a better understanding of the role of different isoforms in innate immune response. Finally, we recommend that alternative splicing quantification should be an integral part of future QTL studies (including single cell studies), and that it will be increasingly relevant to understand the transcriptomic effects of disease-associated risk loci.

## Methods

### Differentiation of Induced Pluripotent Stem Cell (iPSC) lines into macrophages

iPSCs obtained from healthy donors of European descent were selected from the HipSci Consortium^11^, and were differentiated to macrophages using a previously published protocol. Of 315 lines initially selected, 227 (71.6%) were successfully differentiated. RNA-seq libraries were produced for 217, which represented 209 lines after quality control. Differentiated macrophages have been shown to be transcriptionally similar to monocyte-derived macrophages^60^. The differentiated naive macrophages were then stimulated with a diverse panel of adjuvants, resulting in 10 stimulation conditions, as well as naive differentiated (Ctrl_6 and Ctrl_24) and undifferentiated controls (Prec_D0 and Prec_D2). mRNA was harvested 6 and 24 hours after stimulation. Multiple cell lines were derived from a varying number of individuals in each stimulation condition (ranging between 177-202 individuals), resulting in a total of 4,698 unique RNA-seq libraries across all conditions (Panousis et al. 2023; in press). Detailed experimental protocol for this experiment is provided by Panousis et al. 2023 (in press).

### Genotype imputation and quality control

Individuals who donated cell lines were previously genotyped through the HipSci project^11^. Genotypes were obtained for each cell line from from the HipSci Consortium (hipsci.org) and genotype calling and imputation to UK10K and 1000 Genomes Phase I is described in ref 11. We used CrossMap^65^to lift over from GRCh37 to GRCh38. Imputed variants with a low imputation score (INFO <0.4), Hardy-Weinberg equilibrium P-value < 10^-6^, a minor allele frequency (MAF) < 0.05 or a missingness rate > 0.001 were filtered out. For the remaining variants, genotype principal components (PCs) were calculated using EIGENSTRAT^66^ to correct for population stratification.

### RNA-seq quality control and read mapping

RNA-seq reads (FASTQ files) were mapped to the reference genome build GRCh38 using splice-aware aligner STAR v2.6.1^67^using the following parameters:

“--twopassMode Basic --outSAMstrandField intronMotif --outSAMtype BAM SortedByCoordinate --outSAMunmapped Within --outSAMattributes All -- outFilterMismatchNoverReadLmax 0.04 --outSAMmultNmax 1 --limitBAMsortRAM 40000000000 --sjdbOverhang 74 --waspOutputMode SAMtag”

This step outputs BAM files that were then used for the identification of split reads using LeafCutter.

### WASP filtering of ambiguously-mapped reads

It has been shown that ambiguously-mapped reads (RNA-seq reads that map to multiple genomic positions) can bias the number of split reads mapped across splice junctions^68^. WASP^68^ identifies an ambiguously-mapped read by replacing variant positions with the alternative alleles and then remapping. In the previous step, we used STAR with the “waspOutputMode” which flags reads that pass the wasp filter with “[vW]=1”. We therefore only removed read alignments which did not have tag “[-vW]=1” using this command in SAMtools^69^:

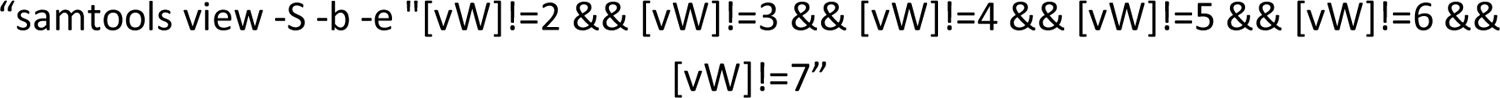

### Identification of split reads

Split reads were identified from BAM files from the previous step using regtools^70^ and output as .junc files. The following command and parameters were used:

“regtools junctions extract -s 0 -a 8 -m 50 -M 500000”

These parameters specify the minimum and maximum intron length (50 bp and 500,000 bps in length, respectively), and a minimum splice junction anchor length of 8 bps. The last parameter means that there must be reads supporting at least 8 basepairs of either side of a splice junction.

### Intron clustering

Mapped split reads (as .junc files) were used to perform intron clustering using the leafcutter_cluster.py script. In this step, .junc files from all samples within each of the 24 stimulation were generated using this command:

“python leafcutter_cluster_regtools_py3.py -j junc_file -m 50 -l 500000”

A minimum number of 50 split reads across samples was required to support an intron cluster. Maximum intron length used was 500 kbp.

For each identified intron cluster with introns 1…j, LeafCutter quantifies intron usage ratio R for intron k as:

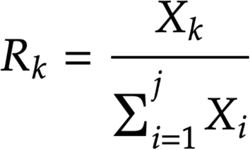

Where X_k_ is the number of split reads that belong to an identified intron k. Differential splicing analysis

Differential splicing analysis was performed between each of the 10 stimulated conditions and their corresponding timepoints controls (e.g. sLPS_6 vs Ctrl_6 and sLPS_24 vs Ctrl_24). LeafCutter differential splicing analysis tool (script leafcutter_ds.R)^12^ was used with eight experimental covariates: run ID, donor, library preparation method, sex, differentiation media, purity, estimated cell diameter, and differentiation time, and with the following parameters:

“–min-samples-per-intron 50 –min-samples-per-group 50 –min-coverage 30 – timeout 900”

### UMAP clustering

UMAP was performed on raw intron usage ratios for introns that were identified across all 24 conditions. UMAP clustering was performed after 3 pre-processing steps. First, we performed quantile normalisation on raw intron usage rations. Second, we performed rank-based inverse normal transformation. Third, we regressed out eight technical covariates similar to the differential splicing analysis: run ID, Donor, library preparation method, sex, differentiation media, purity, estimated cell diameter, and differentiation time. We used the umap R package to perform UMAP clustering (https://github.com/tkonopka/umap).

### Intron usage ratio quality control and normalization

In order to use intron usage as quantitative traits, we first used the LeafCutter script “prepare_phenotype_table.py”, which applies two default filters at the intron-level. It removes introns which have an intron usage ratio of 0 in more than 40% of samples, and those with a standard deviation of less than 5×10^-3^. The script also applies quantile normalization to the introns. These introns were mapped to known intron-exon junctions in genes using the GENCODE v27 annotation in order to map its corresponding gene.

### Mapping sQTLs using intron usage ratios

sQTLs were mapped using normalised intron usage ratios and genotype data from samples within each stimulation condition. Variants in a 1 mega base pair (Mb) window around the transcription start site (TSS) were associated with the intron usage ratios. Genotype-intron association was modelled using a linear regression model implemented in QTLtools^71^ with the parameters:

“-seed 1354145 --permute 1000 --window 1000000 --grp-best”

The option –grep-best allows phenotypes (intron usage ratios) to be organized in groups. Within each phenotype group, the genotype-phenotype sample labels are permuted in exactly the same way. This allows P-values for phenotypes within the same group to be comparable and for QTLtools to report the best associated phenotype per group. We group introns by the gene they belong to, and report the best associated variant-intron per gene.

In all of our analyses, we included the first three genotypic principal components (PC) as covariates. In order to remove unwanted variability in intron usage ratios, we mapped sQTLs separately in different conditions using 0, 1, 2, 3, 4, 5, 6, 7, 8, 9, 10, 20 and 50 intron usage ratio PCs as additional covariates. We then counted the number of genes with significant sQTL effects (sGene) at a false discovery rate (FDR) <= 0.05 using the R package qvalue v2.16.0^72^. In all our downstream analyses, we use the number of intron usage ratio PCs that maximises the number of sGenes discovered.

### Genome-wide summary statistics

GWAS summary statistics were downloaded from either the GWAS catalogue^73^ or from UK BioBank GWAS summary statistics^74^ (Supplementary Table S2). Summary statistics were formatted using a custom script so that each file contains at least: the chromosome and position (GRCh38) of each associated variant, effect size, and standard error.

### Identification of genome-wide significant loci from IMD GWAS summary statistics and colocalisation analysis

To identify genome-wide significant loci, we identified all variants that passed a P-value threshold < 5×10^-8^. We ordered these SNPs from lowest to highest P-value and then iterated over each of them. At each iteration, we define a 1 Mbp window centred around the SNP. If a SNP falls within a locus that has been defined in a previous iteration, it is not used to define a new locus.

For each genome-wide significant locus, we first identified the set of all genes whose TSS falls within the locus. We performed pairwise colocalisation tests for all the intron usage ratios (within each of these genes) and the GWAS association signal. We perform these tests within each of the 24 conditions.

We matched variants between the sQTL and GWAS association signals by position on GRCh38 only since not all GWAS studies provided alleles. We perform colocalisation analysis using the R package coloc v5.1.1^31^ and we discard GWAS-intron pairs that have < 2 variants in common. Only SNPs in common between each GWAS study and our genotypes are considered. This output a list of tested introns per GWAS per condition. The same procedure was used for GTEx sQTLs.

Locus plots for the colocalisation were produced using the locuscomparer R package^75^. Heatmaps for visualising PP4 values were generated using the ComplexHeatmap R packages^76^.

### Condition-specificity analysis using mash

We tested the condition-specificity of sQTLs using mashR v0.2.57^24^. mashR is an adaptive shrinkage framework that can be used to compare effect size estimates in a multi-tissue or multi-condition association study. It outputs shrunk effect size estimates in addition to a statistic indicating if effect sizes between two conditions are significantly different from each other (local false sign rate; LFSR). We use this framework to test if a given sQTL effect is a “response sQTL”, meaning that its effect size is significantly different in a stimulated macrophage condition from unstimulated macrophages.

mashR requires training data to learn the correlation structure in the data from a set of canonical and data-driven covariance matrices (i.e. adaptive). The data-driven matrices are usually derived from the strongest QTL associations in the data. mashR then uses a randomly sampled set of QTL association to learn the mixture component of the provided covariance matrices. We use the lead sQTL SNP per gene (for genes with an FDR < 0.05) as our strong subset, and 10^6^ randomly sampled sQTL associations as the random subset (effect sizes and standard errors). Mash can also be trained using a baseline condition, against which other conditions are compared (we use Ctrl_24 as a baseline condition).

The learned model can be used to recalculate “posterior” summary statistics for any desired set of sQTL associations by providing their effect sizes and standard errors. In our analysis, we recomputed effect sizes and LFSR for two sets of sQTL effects:

1. Lead SNP for significant sQTLs (FDR < 0.05) per sGene within each stimulated condition (Figure 3b). This was used to estimate the number of sGenes with at least one response sQTL.
2. Lead SNP for colocalised genes (PP4 ≥ 0.75) in the condition in which the colocalisation was detected. This was used to estimate the proportion of colocalised genes with a response sQTL within each stimulated condition (Figure 4b).

### Colocalisation between significant sQTLs and eQTLs

We performed statistical colocalisation between all significant sQTLs (FDR < 0.05) to investigate whether the sQTL and eQTL signals in our sGenes are likely to be driven by the same causal variant. We only colocalised the intron with the highest P-value per sGene.

## Acknowledgements

We would like to thank the Human Genetics Informatics team at the Wellcome Sanger Institute for providing computational support to run the analyses described in this manuscript. This work was supported by Wellcome Sanger Institute Core funding from the Wellcome Trust (206194, 220540/Z/20/A). The iPSC lines were generated at the Wellcome Sanger Institute, under the Human Induced Pluripotent Stem Cell Initiative funded by a strategic award (WT098503) from the Wellcome Trust and Medical Research Council. NIP was supported by the Early Postdoc Mobility fellowship from the Swiss National Science Foundation. We would like to thank Emma Davenport and Alex Tokolyi for reviewing this manuscript and providing valuable feedback.

## Author Contributions

OEG performed the analyses and, along with NIP, CAA and DJG drafted the manuscript with contributions from all authors. NIP assisted with multiple analyses and independently validated sQTL results and provided feedback to improve the manuscript. NK provided statistical feedback on several occasions, assisted with multiple analyses, and conducted analysis on the pilot data. MI, LBV, AT and CG applied the differentiation protocol, performed QC metrics on the differentiated macrophages and carried out the stimulations under the supervision of CG. AK optimised the low-input bulk RNA-seq protocol and prepared the RNA-seq libraries along with MI and AB. DJG conceptualised the study while CAA and DJC supervised this work.

## Data availability

Imputed genotype data for the HipSci lines, unprocessed RNA-seq data are available, and processed read counts (TPMs and splice junction usage ratios) will be available on ENA and EGA (accession numbers available upon publication). Full sQTL summary statistics are available on Synapse (DOI: 10.7303/syn51514197).

## References

[1] Maurano, M.T. et al. (2012) ‘Systematic localization of common disease-associated variation in regulatory DNA’, Science, 337(6099), pp. 1190–1195. doi:10.1126/science.1222794.

[2] Schaub, M.A. et al. (2012) ‘Linking disease associations with regulatory information in the human genome’, Genome Research, 22(9), pp. 1748–1759. doi:10.1101/gr.136127.111.

[3] Võsa, U. et al. (2021) ‘Large-scale cis- and Trans-eqtl analyses identify thousands of genetic loci and polygenic scores that regulate blood gene expression’, Nature Genetics, 53(9), pp. 1300–1310. doi:10.1038/s41588-021-00913-z.

[4] Aguet, F. et al. (2020) ‘The GTEX consortium atlas of genetic regulatory effects across human tissues’, Science, 369(6509), pp. 1318–1330. doi:10.1126/science.aaz1776.

[5] Kerimov, N. et al. (2021) ‘A compendium of uniformly processed human gene expression and splicing quantitative trait loci’, Nature Genetics, 53(9), pp. 1290–1299. doi:10.1038/s41588-021-00924-w.

[6] Chun, S. et al. (2017) ‘Limited statistical evidence for shared genetic effects of eqtls and autoimmune-disease-associated loci in three major immune-cell types’, Nature Genetics, 49(4), pp. 600–605. doi:10.1038/ng.3795.

[7] Mostafavi, H. et al. (2022) Limited overlap of eqtls and GWAS hits due to systematic differences in Discovery [Preprint]. doi:10.1101/2022.05.07.491045.

[8] Li, Y.I. et al. (2016) ‘RNA splicing is a primary link between genetic variation and disease’, Science, 352(6285), pp. 600–604. doi:10.1126/science.aad9417.

[9] Kim-Hellmuth, S. et al. (2020) ‘Cell type–specific genetic regulation of gene expression across human tissues’, Science, 369(6509). doi:10.1126/science.aaz8528.

[10] Kilpinen, H. et al. (2017) ‘Common genetic variation drives molecular heterogeneity in human ipscs’, Nature, 546(7658), pp. 370–375. doi:10.1038/nature22403.

[11] Li, Y.I. et al. (2017) ‘Annotation-free quantification of RNA splicing using leafcutter’, Nature Genetics, 50(1), pp. 151–158. doi:10.1038/s41588-017-0004-9.

[12] Liu, H. et al. (2018) ‘Alternative splicing analysis in human monocytes and macrophages reveals MBNL1 as major regulator’, Nucleic Acids Research, 46(12), pp. 6069–6086. doi:10.1093/nar/gky401.

[13] Green, I.D. et al. (2020) ‘Macrophage development and activation involve coordinated intron retention in key inflammatory regulators’, Nucleic Acids Research, 48(12), pp. 6513– 6529. doi:10.1093/nar/gkaa435.

[14] Stenmark, H. (2009) ‘Rab GTPases as coordinators of vesicle traffic’, Nature Reviews Molecular Cell Biology, 10(8), pp. 513–525. doi:10.1038/nrm2728.

[15] Yoneyama, M. et al. (2005) ‘Shared and unique functions of the dexd/h-box helicases rig-I, MDA5, and LGP2 in antiviral innate immunity’, The Journal of Immunology, 175(5), pp. 2851–2858. doi:10.4049/jimmunol.175.5.2851.

[16] Kato, H. et al. (2005) ‘Cell type-specific involvement of rig-I in antiviral response’, Immunity, 23(1), pp. 19–28. doi:10.1016/j.immuni.2005.04.010.

[17] Castanier, C. et al. (2012) ‘Mavs ubiquitination by the E3 ligase TRIM25 and degradation by the proteasome is involved in type I interferon production after activation of the antiviral rig-i-like receptors’, BMC Biology, 10(1). doi:10.1186/1741-7007-10-44.

[18] AL Hamrashdi, M. and Brady, G. (2022) ‘Regulation of IRF3 activation in human antiviral signaling pathways’, Biochemical Pharmacology, 200, p. 115026. doi:10.1016/j.bcp.2022.115026.

[19] Saleh, D. et al. (2017) ‘Kinase activities of RIPK1 and RIPK3 can direct IFN-β synthesis induced by lipopolysaccharide’, The Journal of Immunology, 198(11), pp. 4435–4447. doi:10.4049/jimmunol.1601717.

[20] Gack, M.U. et al. (2009) ‘Influenza A virus NS1 targets the ubiquitin ligase TRIM25 to evade recognition by the host viral RNA sensor rig-I’, Cell Host & Microbe, 5(5), pp. 439–449. doi:10.1016/j.chom.2009.04.006.

[21] Liao, K.-C. and Garcia-Blanco, M.A. (2021) ‘Role of alternative splicing in regulating host response to viral infection’, Cells, 10(7), p. 1720. doi:10.3390/cells10071720.

[22] Lad, S.P. et al. (2008) ‘Identification of mavs splicing variants that interfere with Rigi/MAVS pathway signaling’, Molecular Immunology, 45(8), pp. 2277–2287. doi:10.1016/j.molimm.2007.11.018.

[23] Wagner, A.R. et al. (2021) ‘Global transcriptomics uncovers distinct contributions from splicing regulatory proteins to the macrophage innate immune response’, Frontiers in Immunology, 12. doi:10.3389/fimmu.2021.656885.

[24] Urbut, S.M. et al. (2018) ‘Flexible statistical methods for estimating and testing effects in genomic studies with multiple conditions’, Nature Genetics, 51(1), pp. 187–195. doi:10.1038/s41588-018-0268-8.

[25] Unuvar Purcu D, Korkmaz A, Gunalp S, Helvaci DG, Erdal Y, Dogan Y, et al. (2022) Effect of stimulation time on the expression of human macrophage polarization markers. PLoS ONE 17(3): e0265196. https://doi.org/10.1371/journal.pone.0265196

[26] Sharif, O., Bolshakov, V.N., Raines, S. et al. Transcriptional profiling of the LPS induced NF-κB response in macrophages. BMC Immunol 8, 1 (2007). https://doi.org/10.1186/1471-2172-8-1

[27] Garrido-Martín, D., Borsari, B., Calvo, M. et al. Identification and analysis of splicing quantitative trait loci across multiple tissues in the human genome. Nat Commun 12, 727 (2021). https://doi.org/10.1038/s41467-020-20578-2

[28] Wang, E., Sandberg, R., Luo, S. et al. Alternative isoform regulation in human tissue transcriptomes. Nature 456, 470–476 (2008). https://doi.org/10.1038/nature07509

[29] Lappalainen, T., Sammeth, M., Friedländer, M. et al. Transcriptome and genome sequencing uncovers functional variation in humans. Nature 501, 506–511 (2013). https://doi.org/10.1038/nature12531

[30] Giambartolomei C, Vukcevic D, Schadt EE, Franke L, Hingorani AD, Wallace C, et al. (2014) Bayesian Test for Colocalisation between Pairs of Genetic Association Studies Using Summary Statistics. PLoS Genet 10(5): e1004383. https://doi.org/10.1371/journal.pgen.1004383

[31] Annalisa Buniello, Jacqueline A L MacArthur, Maria Cerezo, Laura W Harris, James Hayhurst, Cinzia Malangone, Aoife McMahon, Joannella Morales, Edward Mountjoy, Elliot Sollis, Daniel Suveges, Olga Vrousgou, Patricia L Whetzel, Ridwan Amode, Jose A Guillen, Harpreet S Riat, Stephen J Trevanion, Peggy Hall, Heather Junkins, Paul Flicek, Tony Burdett, Lucia A Hindorff, Fiona Cunningham, Helen Parkinson, The NHGRI-EBI GWAS Catalog of published genome-wide association studies, targeted arrays and summary statistics 2019, *Nucleic Acids Research*, Volume 47, Issue D1, 08 January 2019, Pages D1005–D1012, https://doi.org/10.1093/nar/gky1120

[32] Mountjoy, E. et al. (2021) ‘An open approach to systematically prioritize causal variants and genes at all published human GWAS trait-associated loci’, Nature Genetics, 53(11), pp. 1527–1533. doi:10.1038/s41588-021-00945-5.

[33] Symoens S, Malfait F, Vlummens P, Hermanns-Lê T, Syx D, De Paepe A (2011) A Novel Splice Variant in the N-propeptide of *COL5A1* Causes an EDS Phenotype with Severe Kyphoscoliosis and Eye Involvement. PLoS ONE 6(5): e20121. https://doi.org/10.1371/journal.pone.0020121

[34] Anna, A., Monika, G. Splicing mutations in human genetic disorders: examples, detection, and confirmation. J Appl Genetics 59, 253–268 (2018). https://doi.org/10.1007/s13353-018-0444-7

[35] Sanz DJ, Hollywood JA, Scallan MF, Harrison PT (2017) Cas9/gRNA targeted excision of cystic fibrosis-causing deep-intronic splicing mutations restores normal splicing of *CFTR* mRNA. PLoS ONE 12(9): e0184009. https://doi.org/10.1371/journal.pone.0184009

[36] Nellore, A., Jaffe, A.E., Fortin, JP. et al. Human splicing diversity and the extent of unannotated splice junctions across human RNA-seq samples on the Sequence Read Archive. Genome Biol 17, 266 (2016). https://doi.org/10.1186/s13059-016-1118-6

[37] Pickrell JK, Pai AA, Gilad Y, Pritchard JK (2010) Noisy Splicing Drives mRNA Isoform Diversity in Human Cells. PLoS Genet 6(12): e1001236. https://doi.org/10.1371/journal.pgen.1001236

[38] Parlato M, Nian Q, Charbit-Henrion F, Ruemmele FM, Rodrigues-Lima F, Cerf-Bensussan N; Immunobiota Study Group. Loss-of-Function Mutation in PTPN2 Causes Aberrant Activation of JAK Signaling Via STAT and Very Early Onset Intestinal Inflammation. Gastroenterology. 2020 Nov;159(5):1968-1971.e4. doi: 10.1053/j.gastro.2020.07.040. Epub 2020 Jul 25. PMID: 32721438.

[39] Pike KA and Tremblay ML (2018) Protein Tyrosine Phosphatases: Regulators of CD4 T Cells in Inflammatory Bowel Disease. Front. Immunol. 9:2504. doi: 10.3389/fimmu.2018.02504

[40] Spalinger, M.R. et al. (2018) ‘PTPN2 regulates inflammasome activation and controls onset of intestinal inflammation and colon cancer’, Cell Reports, 22(7), pp. 1835–1848. doi:10.1016/j.celrep.2018.01.052.

[41] Spalinger, M.R. et al. (2022) ‘Loss of protein tyrosine phosphatase non-receptor type 2 reduces il-4-driven alternative macrophage activation’, Mucosal Immunology, 15(1), pp. 74–83. doi:10.1038/s41385-021-00441-3.

[42] Sigismund, S., Lanzetti, L., Scita, G. et al. Endocytosis in the context-dependent regulation of individual and collective cell properties. Nat Rev Mol Cell Biol 22, 625–643 (2021). https://doi.org/10.1038/s41580-021-00375-5

[43] Martin Steger, Francesca Tonelli, Genta Ito, Paul Davies, Matthias Trost, Melanie Vetter, Stefanie Wachter, Esben Lorentzen, Graham Duddy, Stephen Wilson, Marco AS Baptista, Brian K Fiske, Matthew J Fell, John A Morrow, Alastair D Reith, Dario R Alessi, Matthias Mann (2016) Phosphoproteomics reveals that Parkinson’s disease kinase LRRK2 regulates a subset of Rab GTPases eLife 5:e12813 https://doi.org/10.7554/eLife.12813

[44] Bae, EJ., Kim, DK., Kim, C. et al. LRRK2 kinase regulates α-synuclein propagation via RAB35 phosphorylation. Nat Commun 9, 3465 (2018). https://doi.org/10.1038/s41467-018-05958-z

[45] Luis Bonet-Ponce et al. LRRK2 mediates tubulation and vesicle sorting from lysosomes.Sci. Adv.6,eabb2454(2020).DOI:10.1126/sciadv.abb2454

[46] Bae, EJ., Kim, DK., Kim, C. et al. LRRK2 kinase regulates α-synuclein propagation via RAB35 phosphorylation. Nat Commun 9, 3465 (2018). https://doi.org/10.1038/s41467-018-<otherinfo>05958-z</otherinfo>

[47] Lee, H.-S. et al. (2020) ‘Inflammatory bowel disease and parkinson’s disease: Common pathophysiological links’, Gut [Preprint]. doi:10.1136/gutjnl-2020-322429.

[48] Mrozowska, P.S. and Fukuda, M. (2016) ‘Regulation of podocalyxin trafficking by Rab small GTPases in epithelial cells’, Small GTPases, 7(4), pp. 231–238. doi:10.1080/21541248.2016.1211068.

[49] Kinoshita, R., Homma, Y. and Fukuda, M. (2020) ‘Rab35–gefs, DENND1A and folliculin differentially regulate podocalyxin trafficking in two- and three-dimensional epithelial cell cultures’, Journal of Biological Chemistry, 295(11), pp. 3652–3663. doi:10.1074/jbc.ra119.011646.

[50] Mu, Z., Wei, W., Fair, B. et al. The impact of cell type and context-dependent regulatory variants on human immune traits. Genome Biol 22, 122 (2021). https://doi.org/10.1186/s13059-021-02334-x

[51] Eugene Melamud, John Moult, Structural implication of splicing stochastics, *Nucleic Acids Research*, Volume 37, Issue 14, 1 August 2009, Pages 4862–4872, https://doi.org/10.1093/nar/gkp444

[52] Wright, D.J., Hall, N.A.L., Irish, N. et al. Long read sequencing reveals novel isoforms and insights into splicing regulation during cell state changes. BMC Genomics 23, 42 (2022). https://doi.org/10.1186/s12864-021-08261-2

[53] Rotival, M., Quach, H. & Quintana-Murci, L. Defining the genetic and evolutionary architecture of alternative splicing in response to infection. Nat Commun 10, 1671 (2019). https://doi.org/10.1038/s41467-019-09689-7

[54] Mazin, P.V., Khaitovich, P., Cardoso-Moreira, M. et al. Alternative splicing during mammalian organ development. Nat Genet 53, 925–934 (2021). https://doi.org/10.1038/s41588-021-00851-w

[55] Deininger, P. *Alu* elements: know the SINEs. Genome Biol 12, 236 (2011). https://doi.org/10.1186/gb-2011-12-12-236

[56] Jan Attig, Igor Ruiz de los Mozos, Nejc Haberman, Zhen Wang, Warren Emmett, Kathi Zarnack, Julian König, Jernej Ule (2016) Splicing repression allows the gradual emergence of new Alu-exons in primate evolution eLife 5:e19545 https://doi.org/10.7554/eLife.19545

[57] Singer, S.S. et al. (2004) ‘From “junk” to gene: Curriculum vitae of a primate receptor isoform gene’, Journal of Molecular Biology, 341(4), pp. 883–886. doi:10.1016/j.jmb.2004.06.070.

[58] Mercuri, E. et al. (2018) ‘Nusinersen versus sham control in later-onset spinal muscular atrophy’, New England Journal of Medicine, 378(7), pp. 625–635. doi:10.1056/nejmoa1710504.

[59] Roberts, T.C., Langer, R. & Wood, M.J.A. Advances in oligonucleotide drug delivery. Nat Rev Drug Discov 19, 673–694 (2020). https://doi.org/10.1038/s41573-020-0075-7

[60] Alasoo, K., Rodrigues, J., Mukhopadhyay, S. et al. Shared genetic effects on chromatin and gene expression indicate a role for enhancer priming in immune response. Nat Genet 50, 424–431 (2018). https://doi.org/10.1038/s41588-018-0046-7

[61] Lavin, Y. et al. (2014) ‘Tissue-resident macrophage enhancer landscapes are shaped by the local microenvironment’, Cell, 159(6), pp. 1312–1326. doi:10.1016/j.cell.2014.11.018.

[62] Al’Khafaji, A.M. et al. (2021) High-throughput RNA isoform sequencing using programmable cdna concatenation [Preprint]. doi:10.1101/2021.10.01.462818.

[63] Amarasinghe, S.L., Su, S., Dong, X. et al. Opportunities and challenges in long-read sequencing data analysis. Genome Biol 21, 30 (2020). https://doi.org/10.1186/s13059-020-1935-5

[64] Hu, Y., Fang, L., Chen, X. et al. LIQA: long-read isoform quantification and analysis. Genome Biol 22, 182 (2021). https://doi.org/10.1186/s13059-021-02399-8

[65] Hao Zhao and others, CrossMap: a versatile tool for coordinate conversion between genome assemblies, *Bioinformatics*, Volume 30, Issue 7, April 2014, Pages 1006–1007, https://doi.org/10.1093/bioinformatics/btt730

[66] Price, A., Patterson, N., Plenge, R. et al. Principal components analysis corrects for stratification in genome-wide association studies. Nat Genet 38, 904–909 (2006). https://doi.org/10.1038/ng1847

[67] Alexander Dobin and others, STAR: ultrafast universal RNA-seq aligner, *Bioinformatics*, Volume 29, Issue 1, January 2013, Pages 15–21, https://doi.org/10.1093/bioinformatics/bts635

[68] van de Geijn, B., McVicker, G., Gilad, Y. et al. WASP: allele-specific software for robust molecular quantitative trait locus discovery. Nat Methods 12, 1061–1063 (2015). https://doi.org/10.1038/nmeth.3582

[69] Heng Li and others, The Sequence Alignment/Map format and SAMtools, *Bioinformatics*, Volume 25, Issue 16, August 2009, Pages 2078–2079, https://doi.org/10.1093/bioinformatics/btp352

[70] Cotto, K.C., Feng, YY., Ramu, A. et al. Integrated analysis of genomic and transcriptomic data for the discovery of splice-associated variants in cancer. Nat Commun 14, 1589 (2023). https://doi.org/10.1038/s41467-023-37266-6

[71] Delaneau, O., Ongen, H., Brown, A. et al. A complete tool set for molecular QTL discovery and analysis. Nat Commun 8, 15452 (2017). https://doi.org/10.1038/ncomms15452

[72] Storey JD, Bass AJ, Dabney A, Robinson D (2023). qvalue: Q-value estimation for false discovery rate control. R package version 2.32.0, http://github.com/jdstorey/qvalue.

[73] Annalisa Buniello and others, The NHGRI-EBI GWAS Catalog of published genome-wide association studies, targeted arrays and summary statistics 2019, *Nucleic Acids Research*, Volume 47, Issue D1, 08 January 2019, Pages D1005–D1012, https://doi.org/10.1093/nar/gky1120

[74] Zhou, W., Nielsen, J.B., Fritsche, L.G. et al. Efficiently controlling for case-control imbalance and sample relatedness in large-scale genetic association studies. Nat Genet 50, 1335–1341 (2018). https://doi.org/10.1038/s41588-018-0184-y

[75] Boxiang Liu, Michael J. Gloudemans, Abhiram S. Rao, Erik Ingelsson & Stephen B. Montgomery (2019) Abundant associations with gene expression complicate GWAS follow-up, Nature Genetics

[76] Gu, Z. (2016) Complex heatmaps reveal patterns and correlations in multidimensional genomic data. Bioinformatics. DOI: 10.1093/bioinformatics/btw313.

