## Supplementary Figures for "Low-usage splice junctions underpin immune-mediated disease risk"

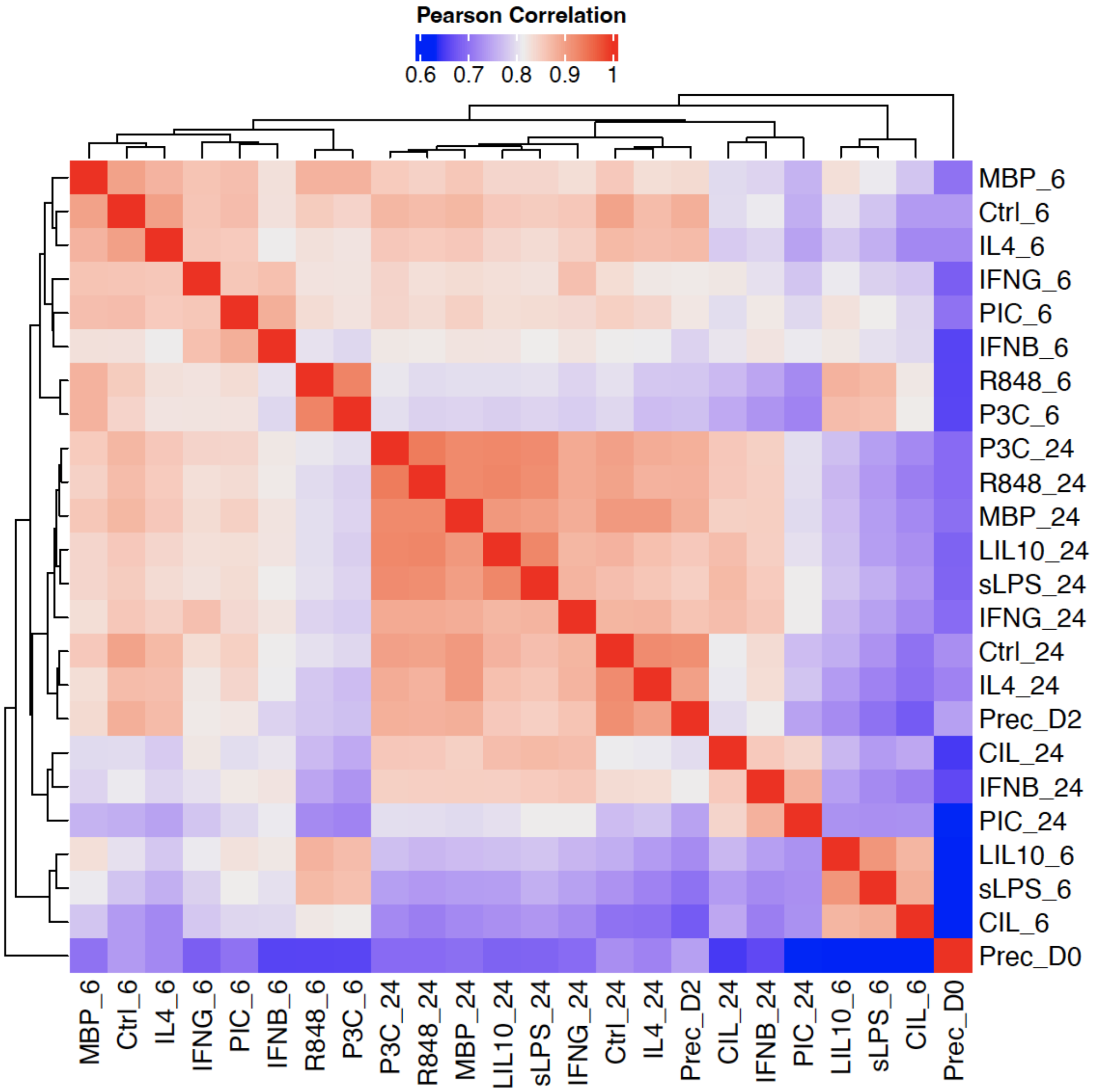


Supplementary Figure S1: Heatmap and dendrogram showing pairwise Pearson correlation coefficients between log-transformed normalised intron usage ratios in introns identified across all conditions. Normalisation consists of regressing out experimental covariates similar to “Differential Splicing Analysis”, quantile normalisation and inverse-normal rank transformation. Only the highest 20,000 variable introns across conditions were used.

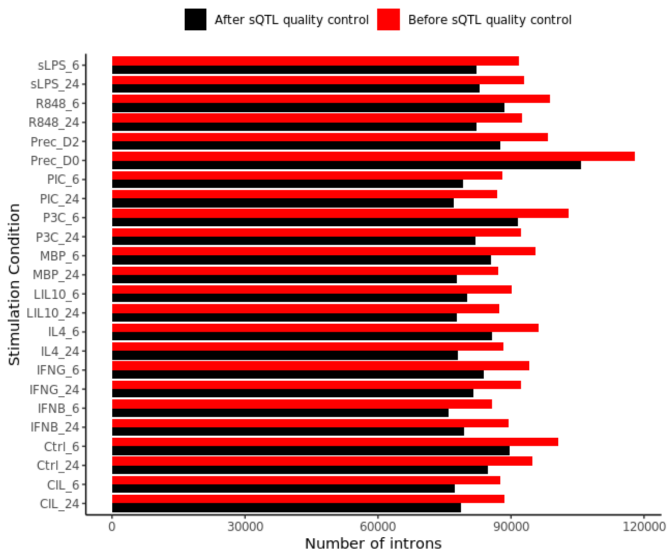


Supplementary Figure S2: Number of introns identified in the intron clustering procedure of leafcutter (before intron sQTL quality control; in red) and the number of introns that were used as input to map sQTLs (after intron sQTL quality control; in black). For quality control steps and parameters see Methods.

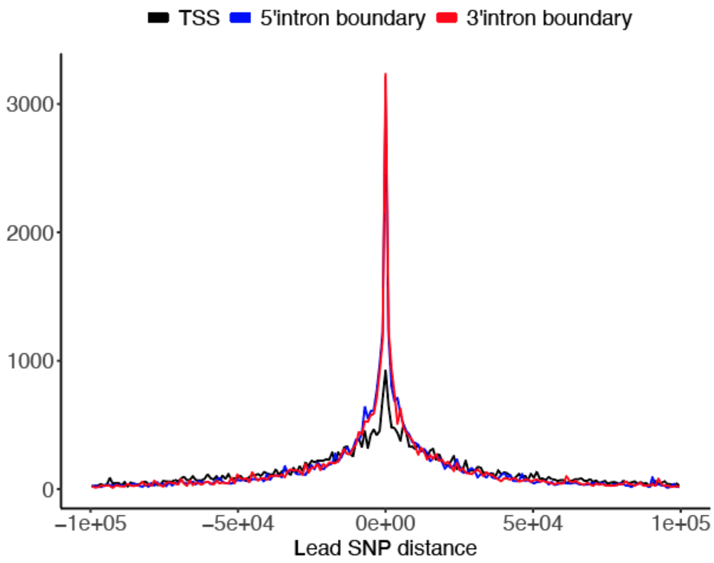


Supplementary Figure S3: Distribution of the distance between the lead SNPs of significant sQTL effects (across all conditions) and transcription start site (TSS; in black) of the sQTL gene, 5’intron boundary (in blue) and 3’ intron boundary (in red).

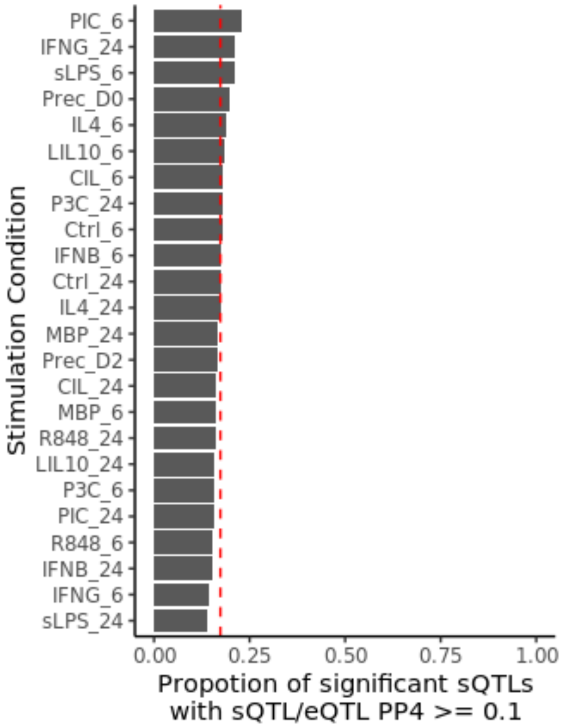


Supplementary Figure S4: Proportion of significant sQTL effects that share a single causal variant (PP4 ≥ 0.1) with the same eQTL gene in the same condition. Red line indicates the average proportion of sQTLs with sQTL/eQTL colocalisation >= 0.1 across conditions.

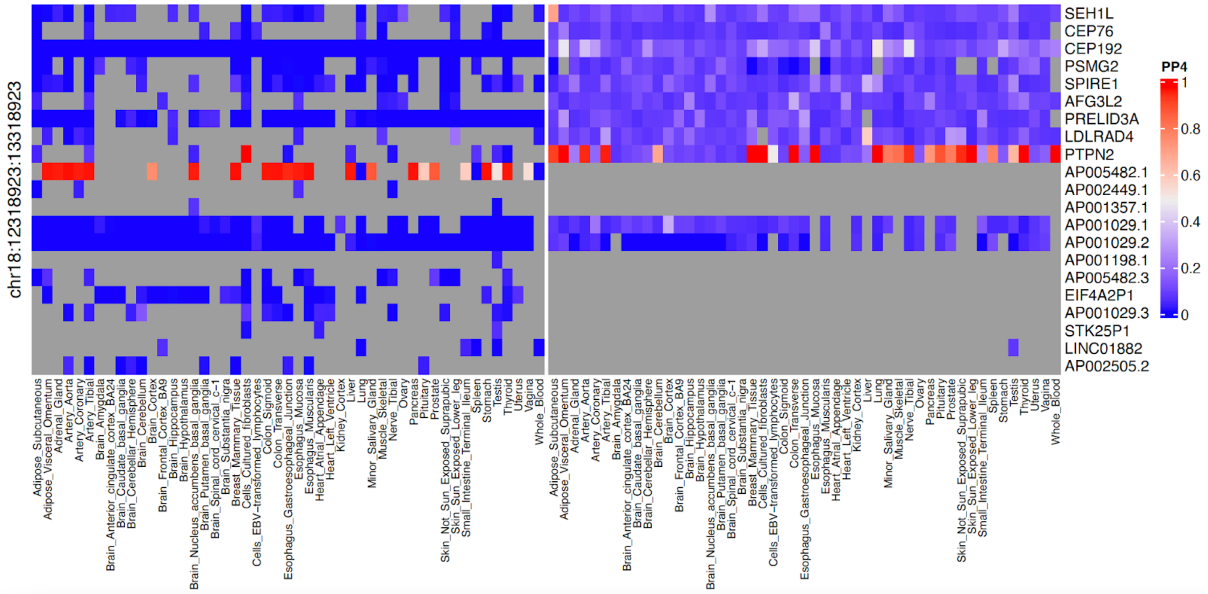


Supplementary Figure S5: Heatmap of PP4 values for all genes in 18p21.11 (rows) and GTEx tissues (columns) split by type of QTL (eQTL on right and sQTL on left).

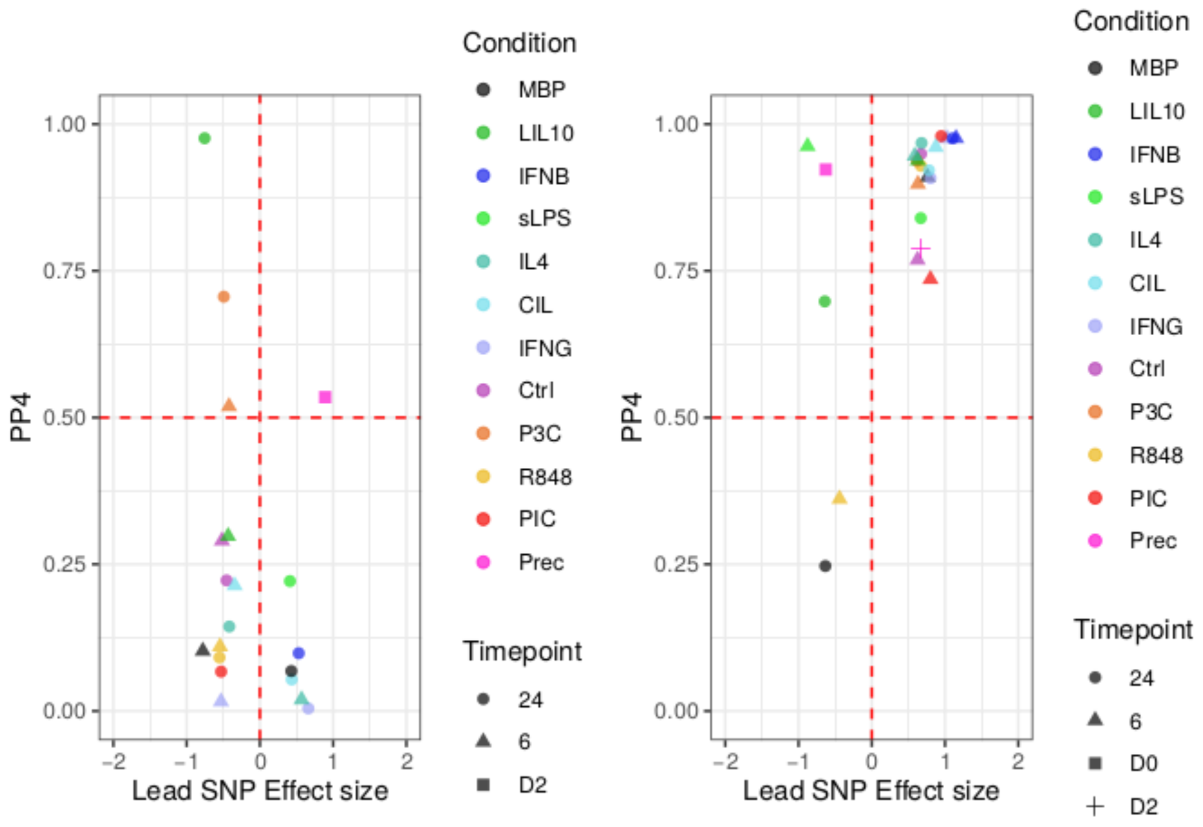


Supplementary Figure S6: effect sizes of the most significant SNP and colocalisation PP4 for the colocalised splice junctions of *LRRK2* (left) and *DENND1B* (right).

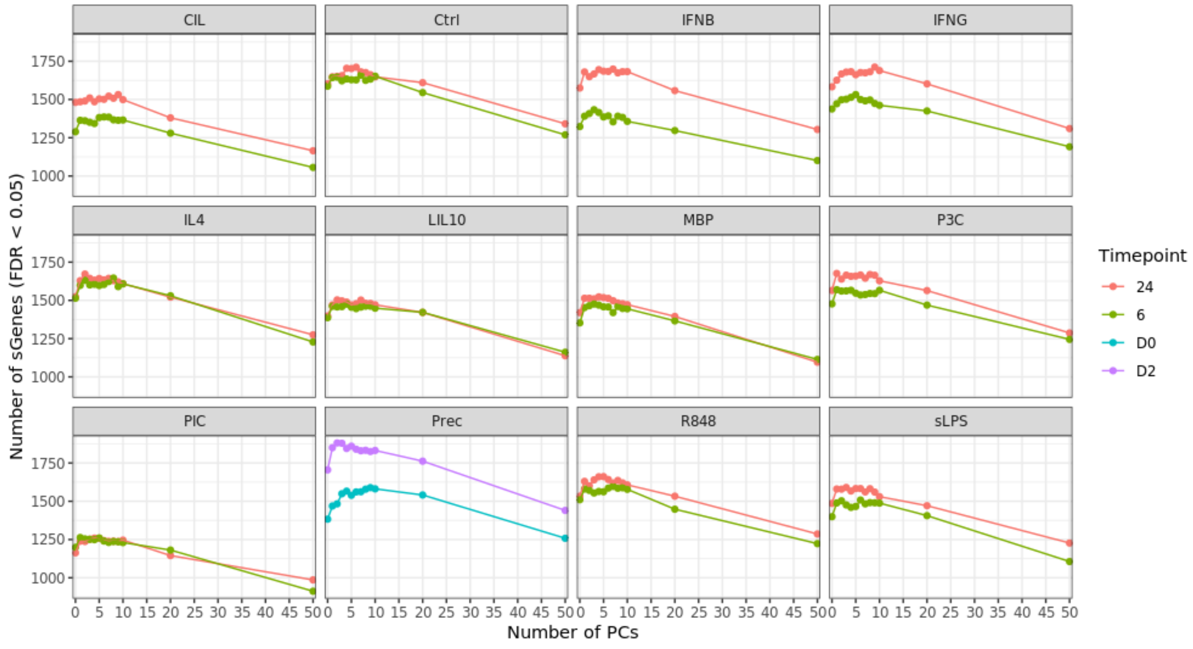

Supplementary Figure S7: Number of intron usage ratio principal components used as covariates (x-axis) versus the number of genes for which a significant sQTL effect was found (y-axis) coloured by time point (6=6 hours, 24=24 hours, D0=Day 0, D2=Day 2).
